# Tumor-secreted versican co-opts myeloid IKKβ during metastasis

**DOI:** 10.1101/2021.05.20.444963

**Authors:** Magda Spella, Anne-Sophie Lamort, Malamati Vreka, Antonia Marazioti, Ioannis Lilis, Giannoula Ntaliarda, Georgios Skiadas, Eleni Bouloukou, Georgia A. Giotopoulou, Mario A.A. Pepe, Stefanie A.I. Weiss, Agnese Petrera, Stefanie M. Hauck, Ina Koch, Michael Lindner, Rudolph A. Hatz, Juergen Behr, Kristina A.M. Arendt, Ioanna Giopanou, David Brunn, Rajkumar Savai, Dieter E. Jenne, Fiona E. Yull, Timothy S. Blackwell, Georgios T. Stathopoulos

## Abstract

The mechanisms tumor cells use to hijack the immune system are largely uncharted. Here we used bioluminescent nuclear factor (NF)-κB reporter mice and macrophages to discover that metastatic tumors trigger NF-κB activation in host macrophages, dependent on mutant *KRAS* signaling and delivered via secretory versican. Versican activates NF-κB in tumor-associated macrophages via inhibitor of NF-κB kinase (IKK) β, resulting in release of interleukin (IL)-1β into the tumor microenvironment. Versican silencing in cancer cells or conditional IKKβ deletion in macrophages prevents myeloid NF-κB activation and metastasis. Versican is overexpressed and/or mutated in human cancers and metastatic effusions with *KRAS* mutations, predicts poor survival, can aid in the development of diagnostic platforms for pleural metastasis, and is druggable via toll-like receptor (TLR) 1/2 inhibition. The data indicate a cardinal role for tumor-derived versican in establishing cross-talk with macrophage IKKβ during metastasis and may foster the development of new therapies and diagnostic tools.

## Introduction

Inflammation is intimately linked with cancer progression and metastasis (1). Oncogenes such as *KRAS* and *MYC* initiate and perpetuate tumor-associated inflammation, and foster proliferation of cancer cells in a cell-autonomous fashion (2-4). In turn, myeloid cells including macrophages, neutrophils, and mast cells, infiltrate human tumors to sustain tumor cell proliferation and angiogenesis (5-7). IL-1β is a major mediator of the protumor effects of cancer-associated myeloid cells and its inhibition via the monoclonal antibody canakinumab was recently shown to possess potentially strong tumor-preventive effects (3, 7-9). Hence, the charting of the signaling mechanisms cancer cells use to recruit and shape the phenotype of myeloid cells in their favor, i.e., to co-opt them, is of paramount importance in the battle to halt tumor metastasis (1). In this regard, oncogenic *KRAS* cooperates with the transcription factor NF-κB in cancer cells to potentiate its protumorigenic functions (3, 10-12). Although NF-κB signaling in cancer cells presents a major tumor promoter, only few studies have addressed the interaction between oncogenes and NF-κB signaling in immune cells (8, 13, 14).

Here we use immunocompetent NF-κB reporter mice and experimental cancer models to identify that *KRAS*-mutant tumors elicit a robust NF-κB response in host macrophages, required for tumor progression and metastasis. We pinned the glycoprotein versican (VCAN) as the tumor-secreted determinant responsible for IKKβ-mediated NF-κB activation, resulting in macrophage differentiation, release of IL-1β into the tumor microenvironment and co-option of macrophages for metastasis. Importantly, the Vcan-NF-κB axis is required for metastasis. VCAN is often mutated in *KRAS-*mutant human cancers and metastatic effusions, and we show that it can be targeted through TLR1/2 inhibition. Our results emphasize a potential clinical role for VCAN, as it can foster the development of a diagnostic platform for pleural metastasis as well as of novel therapeutic tools.

## Results and Discussion

We initiated *in vivo* screens of murine tumor cell lines with known *Kras* mutation status (Figure 1A) (2, 3) by transplanting them into two strains of bioluminescent NF-κB reporter mice expressing ubiquitous *HIV-LTR*.*Luciferase* (*HLL* mice) (15) or *NF-κB*.*GFP*.*Luciferase* (*NGL* mice) (16) transgenes. Pleural injections were selected for tumor cell inoculation because they generate metastatic malignant pleural effusions (MPE) with overt cancer-induced inflammation (2, 3, 7). Serial imaging showed time-dependent NF-κB activation in host cells of recipient mice during pleural metastasis, conditional on the presence of *Kras* mutations in tumor cells (Figure 1,B-E; Supplemental Figure 1,A-B). The NF-κB reporter signal was emitted from pleural tumors and fluid, both containing cancer and immune cells (Figure 1F; Supplemental Figure 1,C-F) (2, 3, 7). Histologic and flow cytometric analysis and quantification localized the NF-κB reporter signal to tumor-infiltrating macrophages of mice with *Kras*-mutant pleural tumors and effusions (Figure 1,G-I; Supplemental Figure 1,G-H; Supplemental Figure 2). Mast cells (7) were not involved in the observed NF-κB response (Supplemental Figure 3,A-B). Time-dependent NF-κB activation in host cells was stronger in MPE-compared with subcutaneous tumor growth-models, and required expression of mutant *Kras* by tumor cells (Supplemental Figure 3,C-I). Adoptive bone marrow transfer corroborated myeloid cells as the origin of tumor-induced NF-κB activation, and pharmacologic killing of pleural macrophages prevented host NF-κB activation and pleural carcinomatosis (Supplemental Figure 4,A-C). The pro-tumor function of pleural macrophages was also consistent with the MPE-protected phenotype of macrophage-depleted *Lyz2*.*Cre;Dta* mice (19) (Supplemental Figure 4,D-E). These data directly show that tumors with *KRAS* mutations can activate NF-κB in macrophages and that these myeloid cells are essential for pleural metastasis.

**Figure 1.**
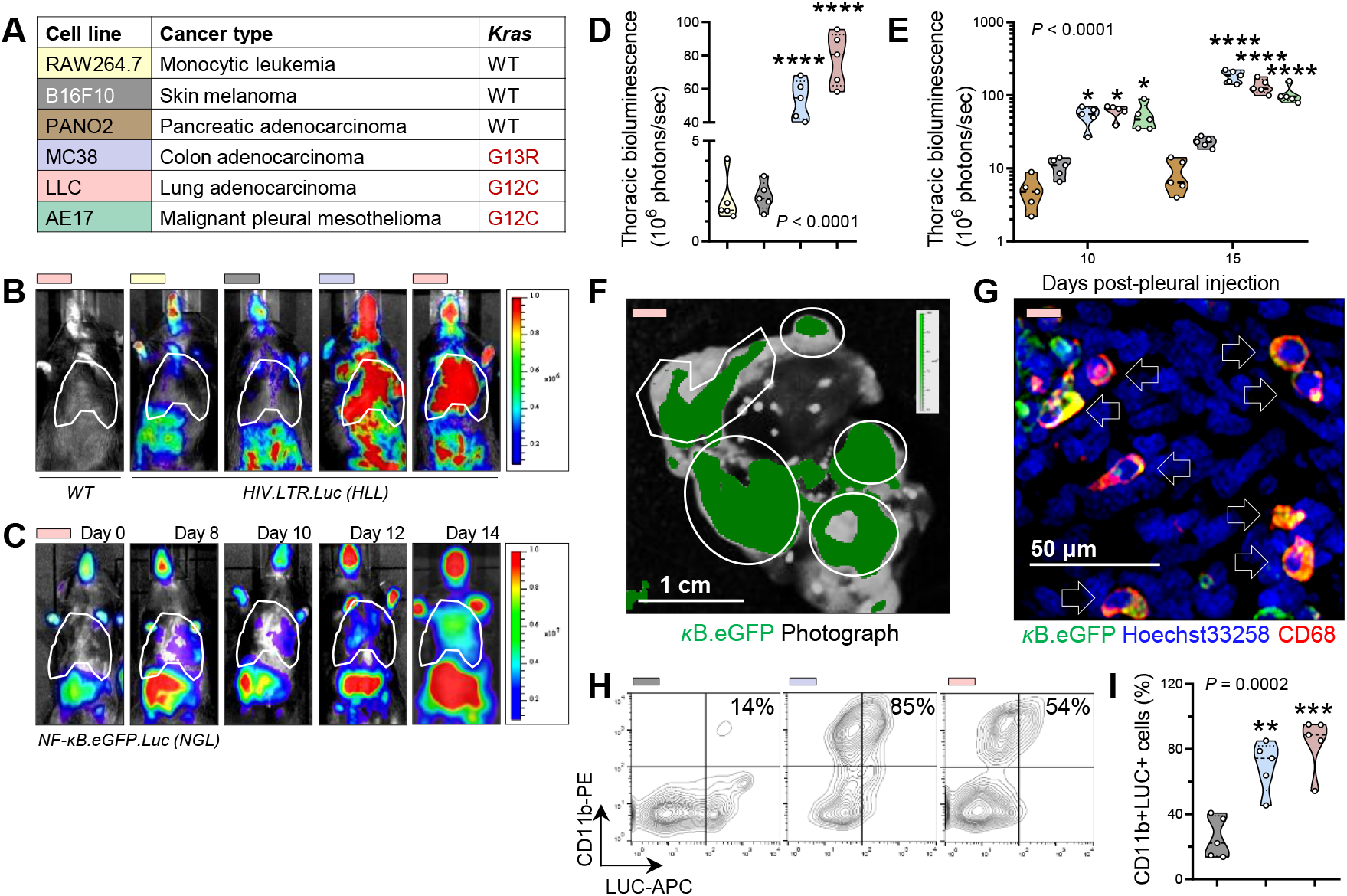
Metastatic *Kras*-mutant tumors activate NF-κB in tumor-infiltrating macrophages. (**A**) Color-coded cancer cells used with *Kras* mutation status. Acronyms are explained in Supplemental Methods. (**B-E**) Bioluminescent images/pseudocolor scale (**B**,**C**) and results (**D**,**E**) from *WT, HLL* (**B**,**D**) (15), and *NGL* (**C**,**E**) (16) NF-κB reporter mice at 14 days (**B**,**D**) or serial time-points (**C**,**E**) post-pleural injection of tumor cells. Bioluminescence is exclusively emitted by host and not tumor cells. (**B**,**C**) Dashed areas delineate the thorax. (**F**) Photographic/biofluorescent image overlay/pseudocolor scale of *NGL* mouse lung explant 14 days post-pleural injection of LLC cells shows NF-κB reporter GFP signal (κB.eGFP) over pleural tumors (outlines; *n* = 10). (**G**) GFP immunoreactivity of pleural tumor sections co-localizes with the macrophage marker CD68 (arrows; *n* = 10). (**H**,**I**) Flow cytometric contour plots (**H**) and data summary (**I**) of pleural tumor cells from *NGL* mice obtained 14 days post-pleural injection stained for the myeloid marker CD11b and the κB.LUC reporter. Percentages in (**H**) pertain to CD11b+LUC+ cells. Data in (**D**,**E**,**I**) are given as raw data (circles), median and quartiles (lines), and kernel density distributions (violin plots) color–coded as in (**A**). Sample size (*n*) =5-10/group; *P*, probability, one- or two-way ANOVA; *, **, ***, and ****, *P* <0.05, *P* <0.01, *P* <0.001, and *P* <0.0001 compared with mice injected with RAW264.7, PANO2, or B16F10 cells at the same time-points, Bonferroni post-tests.

**Figure 2.**
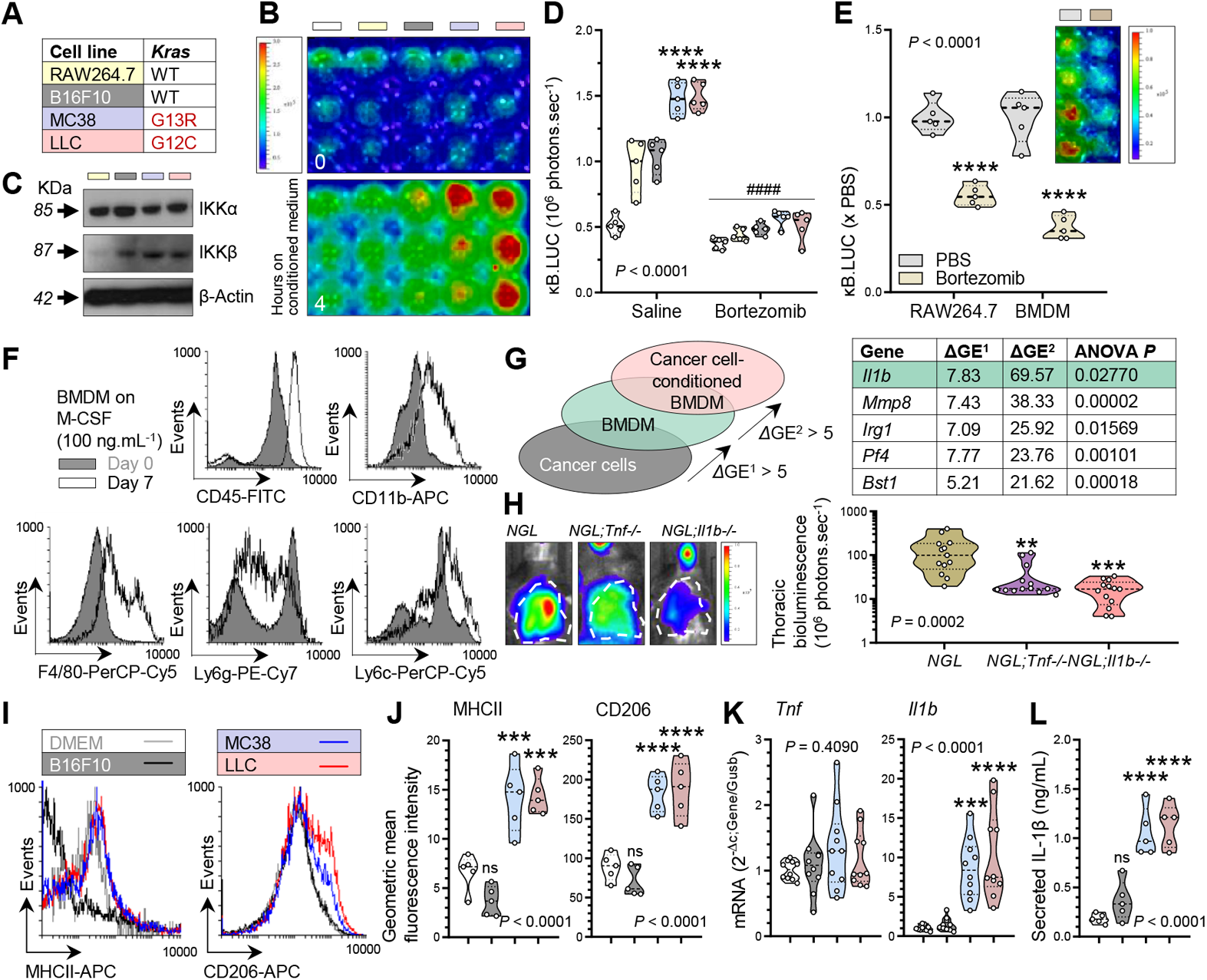
Tumor–secreted mediators drive IKKβ activation, polarization, and IL-1β secretion in macrophages. (**A**) Color-coded cancer cells with *Kras* mutation status. (**B-E**) Bioluminescent images/pseudocolor scale (**B**,**E**), immunoblots (**C**) and results (**D**,**E**) from exposure of RAW264.7 stably expressing p*NGL* (3) (**B-D**) and murine BMDM obtained from *NGL* mice (15) (**E**) to cell-free tumor-conditioned media or DMEM (white boxes) with/without bortezomib pre-treatment (3). (**D**,**E**) *n* =5/group; *P*, probability, one-/two-way ANOVA; **** and ^####^, *P* <0.0001 compared with other groups or non-bortezomib-treated cells, respectively, Bonferroni post-tests. (**F**) Flow cytometric expression histograms of murine bone marrow cells before and after weekly M-CSF exposure. (**G**) Microarray strategy and top-five differentially expressed genes (*Δ*GE) of murine BMDM compared with cancer cells (*Δ*GE^1^) and of tumor-conditioned BMDM compared with naïve BMDM (*Δ*GE^2^). *n* =5/group; *P*, probability, one-way ANOVA. (**H**) Bioluminescent images and data summary of *NGL, NGL;Tnf–/–*, and *NGL;Il1b–/–* mice 14 days post-pleural injection of LLC cells. *n* =13/group; *P*, probability, one-way ANOVA; ** and ***, *P* <0.01 and *P* <0.001 compared with *NGL* mice, Bonferroni post-tests. (**I**-**L**) Histograms (**I**) and results (**J**-**L**) of naïve or tumor-conditioned BMDM for macrophage activation markers (**I**,**J**), *Tnf/Il1b* mRNA (**K**), and IL-1β protein (**L**) expression. *n* =5-10/group; *P*, probability, one-way ANOVA; *** and ****, *P* <0.001 and *P* <0.0001 compared with DMEM and B16F10-conditioned media, Bonferroni post-tests. Data in (**D**,**E**,**H**,**J**,**K**,**L**) are given as raw data (circles), median and quartiles (lines), and kernel density distributions (violin plots) color-coded as in (**A**).

**Figure 3.**
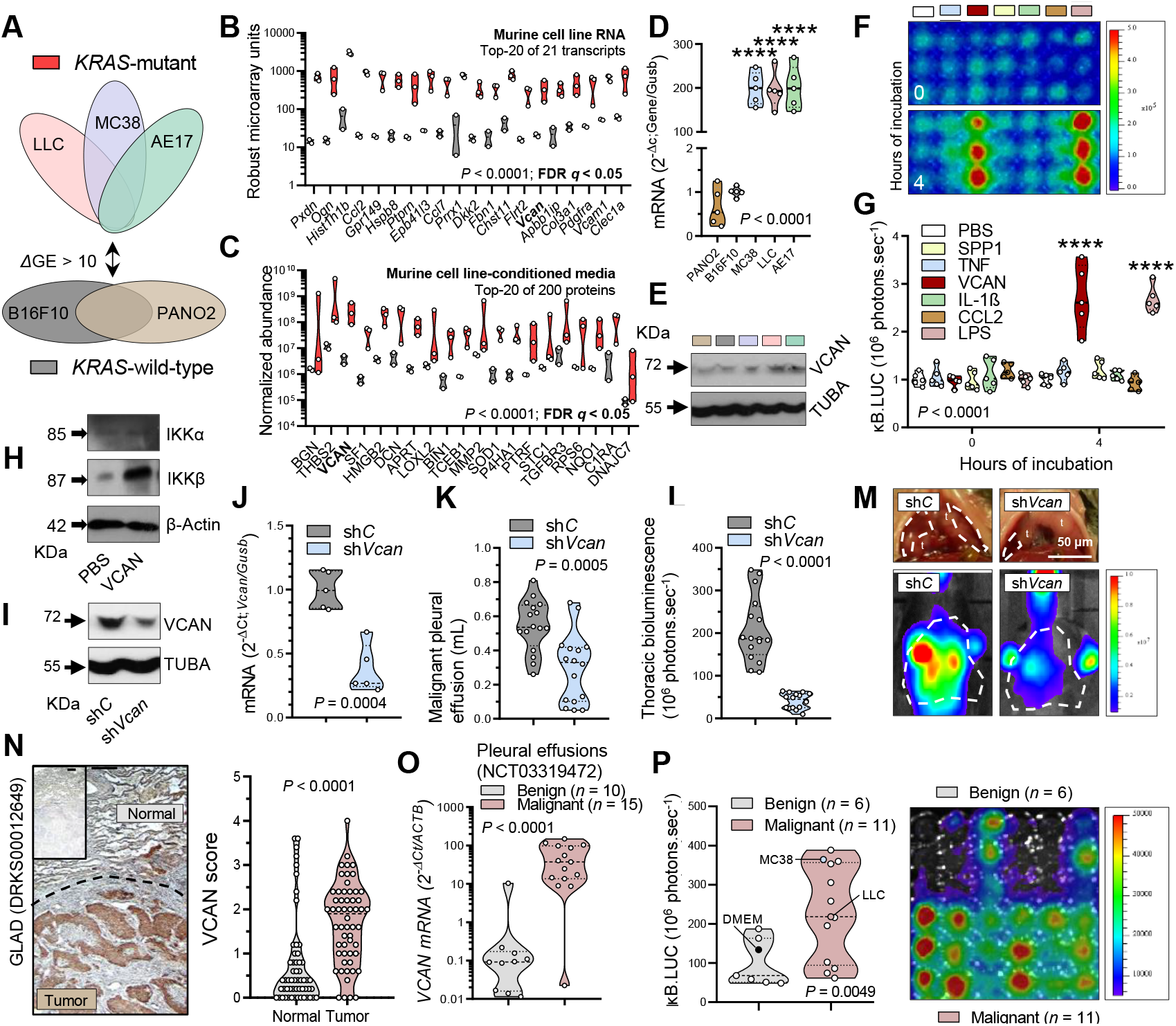
Tumor-secreted versican drives macrophage IKKβ, metastasis, and is a cancer biomarker. (**A-E**) *Kras*-mutant and -wild-type cancer cells were subjected to microarray and LC-MSMS analyses. Experimental design (**A**), top-20 over-represented transcripts (**B**) and secreted proteins (**C**), and *Vcan*/VCAN mRNA/protein expression (**D, E**) of murine cell lines. *n* =2-3/group; *P*, probability, two-way ANOVA; Bold letters, false discovery rate (FDR) *q* <0.05 compared with *Kras*-wild-type cells, two-stage linear step-up procedure (32). (**F**,**G**) Bioluminescent images (**F**) and results (**G**) from p*NGL* RAW264.7 exposed to recombinant proteins. *n* =5/group; *P*, probability, two-way ANOVA; ****, *P* <0.0001 compared with other groups, Bonferroni post-tests. (**H**) Immunoblots of mouse BMDM exposed to VCAN. *n*=5/group. (**I-M**) sh*C* and sh*Vcan* LLC cells were validated and injected intrapleurally into *NGL* mice. Immunoblots (**I**), *Vcan* mRNA expression (**J**), and results (**K**,**L**) and images (**M**) 14 days post-tumor cells. (**I**,**J**) *n* =5/group; (**K-M**) *n* =16/group; *P*, probability, unpaired Student’s t-test. (**N**) Images and VCAN expression of *n* =41 tumor/normal tissue pairs from GLAD study (26). (**O**) *VCAN* mRNA expression of 10 benign and 15 malignant pleural effusions from the MAPED study (25). (**P**) Summary of bioluminescent signal of p*NGL* RAW264.7 after exposure to benign (*n* =6; top triplicates) and malignant (*n* =11; bottom triplicates) pleural effusions and tumor-conditioned media (each triplicate column is one patient). (**N-P**) *P*, probability, unpaired Student’s t-test. (**B-D**,**G**,**J-L**,**N-P**) Shown are raw data (circles), medians/quartiles (lines), and kernel density distributions (violin plots).

**Figure 4.**
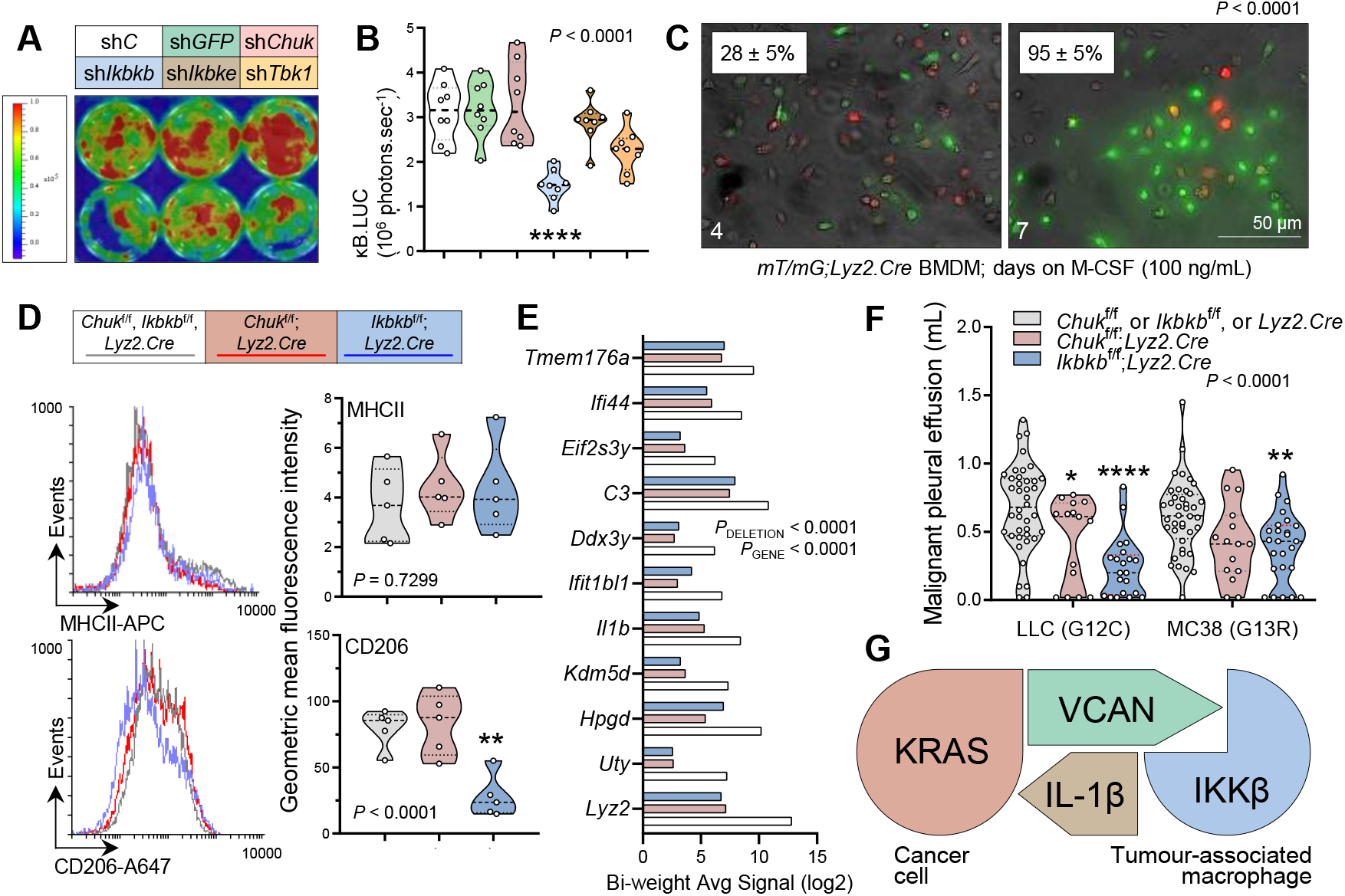
IKKβ mediates NF-κB activity, differentiation, and IL-1β secretion in macrophages to drive metastasis. (**A**,**B**) Bioluminescent image (**A**) and results (**B**) of p*NGL* (3) RAW264.7 72 hours post-infection with sh*C*, sh*GFP*, sh*Chuk*, sh*Ikbkb*, sh*Ikbke*, or sh*Tbk1. n* =8/group; *P*, probability, one-way ANOVA; ****, *P* <0.0001 compared with sh*C*, Bonferroni post-tests. (**C-E**) BMDM were derived from *mT/mG;Lyz2*.*Cre, Chuk*^f/f^*;Lyz2*.*Cre*, and *Ikbkb*^f/f^*;Lyz2*.*Cre* mice. Shown are images and mean ± SD % green cells of bone marrow cells from *mT/mG;Lyz2*.*Cre* mice during/after weekly treatment with M-CSF (**C**), flow cytometric histograms (left) and data summary (right) of marker expression (**D**), and top-differentially expressed genes by microarray (**E**). (**C**,**D**) *n* =5/group; *P*, probability, Fisher’s exact test or one-way ANOVA; **, *P* <0.01 compared with other groups, Bonferroni post-tests. (**E**) *n* =1 pooled triplicate/group; *P*, probabilities, two-way ANOVA. (**F**) *Chuk*^f/f^*;Lyz2*.*Cre* and *Ikbkb*^f/f^*;Lyz2*.*Cre* mice received intrapleural LLC or MC38 cells, and were evaluated after 14 days for MPE. Data summary of *n* =40, 15, and 21 single transgenic control, *Chuk*^f/f^;*Lyz2*.*Cre*, and *Ikbkb*^f/f^;*Lyz2*.*Cre* mice injected with LLC cells, respectively, and of *n* =40, 15, and 25 respective mice injected with MC38 cells. *P*, probability, two-way ANOVA; *, **, and ****, *P* <0.05, *P* <0.01, and *P* <0.0001 compared with controls, Bonferroni post-tests. (**G**) Schematic of the proposed inflammatory loop. *KRAS*-mutant cancer cells secrete VCAN to co-opt IKKβ in macrophages within the metastatic niche, which drives IL-1β secretion by macrophages to foster metastasis.

Tumor-secreted solute factors are responsible for NF-κB activation in macrophages, since murine RAW264.7 macrophages stably expressing the *NGL* plasmid (p*NGL*) (3) responded with robust *in vitro* NF-κB activation to cell-free media conditioned by *Kras*-mutant, but not by *Kras*-wild-type or *Kras*-silenced, tumor cells (Figure 2,A-B; Supplemental Figure 3,H-I). This NF-κB response requires canonical NF-κB signaling, since it involved IKKβ and was attenuated by the proteasome inhibitor bortezomib (Figure 2,C-D; Supplemental Figure 3,H-I; Supplemental Figure 5,A-C). Proteasome-dependent NF-κB activity was also documented in bone marrow-derived macrophages (BMDM) derived from *NGL* mice (Figure 2,E-F). Differential gene expression (*Δ*GE) analyses (GEO datasets GSE94847, GSE94880, GSE130624, and GSE130716; total *n* = 32) identified 13 BMDM-specific transcripts that were further induced by incubation with tumor-conditioned media (*Δ*GE >5; ANOVA *P* <0.05) and included *Il1b* but not *Il6* and *Tnf* reported elsewhere (20) (Figure 2G; Supplemental Tables 1-2). In addition, *NGL* mice deficient in *Il1b* alleles were resistant to tumor-induced NF-κB activation (Figure 2H) and pleural metastasis (3). Incubation of BMDM with *Kras*-mutant tumor-conditioned media promoted their differentiation as assessed by flow cytometry for markers MHCII and CD206, and *Il1b* mRNA and IL-1β protein expression (Figure 2,I-L). These results indicate that solute mediator(s) secreted by *KRAS*-mutant cancer cells during pleural metastasis trigger IKKβ-mediated NF-κB activation, differentiation, and IL-1β elaboration in host macrophages.

We next compared *Kras*-mutant with *Kras*-wild-type cancer cells for secretory molecules triggering macrophage NF-κB activation. Our microarrays analysis identified 25 transcripts over-represented in *Kras*-mutant cells, and a proteomic screen of tumor cell-conditioned media detected 226 proteins secreted >10-fold by *Kras*-mutant over *Kras*-wild-type cells, with *Vcan*/VCAN being common to both screens (Figure 3,A-C; Supplemental Tables 3-5) and validated (Figure 3,D-E; Supplementary Figure 5,D-E). Selected NF-κB ligands were screened using p*NGL*-expressing RAW264.7 cells (Figure 3,F-G), with the TLR2 ligand VCAN potently activating NF-κB in mouse macrophages. VCAN also induced IKKβ in primary murine BMDM, which were verified by microarray to overexpress >10-fold over cancer cells TLR1, TLR2, TLR6-9, and TLR13 (Figure 3H; Supplemental Figure 5,F-H; Supplemental Figure 6; Supplemental Table 6). Importantly, shRNA-mediated *Vcan* silencing in LLC cells diminished their ability to trigger NF-κB activation in *NGL* mice and to precipitate MPE (Figure 3,I-M; Supplemental Figure 5,I-J). Our data indicate that versican secreted by cancer cells triggers IKKβ-mediated NF-κB activation in tumor-associated macrophages and promotes metastasis.

In humans, *VCAN* transcripts were overrepresented in cancers with high *KRAS* mutation frequencies, such as lung adenocarcinomas from smokers (GEO dataset GSE43458) and lung and colorectal cancers (GEO dataset GSE103512) (Supplemental Figure 7,A-B) (21-23). High *VCAN* expression also portended poor survival in a number of other human cancers (http://kmplot.com/analysis/index.php?p=service&cancer=pancancer_rnaseq; Supplemental Figure 7,C-I) (24). Patient samples analysis from two of our clinical studies (Figure 3, N and O), showed that *VCAN* mRNA expression was significantly increased in human MPE compared with benign pleural effusions (BPE) [MAPED study, (25)] and that VCAN protein expression was significantly increased in tumor compared with adjacent lung tissues [GLAD study, (26)]. To test whether the proposed inflammatory loop can serve as a diagnostic tool to distinguish MPE from BPE, which is an unmet clinical need (25), p*NGL*-expressing RAW264.7 macrophages were exposed to cell-free supernatants from human benign and malignant pleural effusions. After 4 hours, a robust NF-κB reporter signal was triggered only by MPE supernatants (Figure 3P). Hence, versican is overexpressed in multiple human cancers, especially in association with *KRAS*-mutations, predicts poor survival, and may be useful as a diagnostic marker of metastatic cancers.

To identify the IKK’s responsible for NF-κB signaling in macrophages, we silenced four such kinases (encoded by the murine *Chuk, Ikbkb, Ikbke*, and *Tbk1* genes) in RAW264.7 macrophages and showed that IKKβ mediates NF-κB activation in these cells (Figure 4,A-B). To further define the functions of IKKβ in macrophages, we obtained BMDM from intercrosses of *Lyz2*.*Cre* mice (19) with mice carrying conditionally-deleted alleles of IKKα (*Chuk*^f/f^) and IKKβ (*Ikbkb*^f/f^) (27-29), as well as with *Cre*-reporter mice switching from red to green fluorescence upon recombination (*mT/mG*) (30). Treatment of bone marrow cells from *mT/mG;Lyz2*.*Cre* mice with M-CSF to drive them towards macrophage differentiation and LYZ2 expression yielded efficient *Cre*-mediated recombination (Figure 4C). Flow cytometric assessment of BMDM derived from control, *Chuk*^f/f^;*Lyz2*.*Cre*, and *Ikbkb*^f/f^;*Lyz2*.*Cre* mice showed that intact IKKβ signaling in primary macrophages is essential for their differentiation and expression of critical proinflammatory genes including *Lyz2, Il1b*, and *C3* (Figure 4,D-E; Supplemental Tables 7-8). Finally, two different syngeneic tumor cell lines, featuring *Kras* mutations and Vcan overexpression, were inoculated into the pleural space of control, *Chuk*^f/f^;*Lyz2*.*Cre*, and *Ikbkb*^f/f^;*Lyz2*.*Cre* mice, to reveal that intact IKKβ signaling in macrophages is required for metastatic MPE (Figure 4F). Thus, VCAN-driven IKKβ activation mediates NF-κB signaling, IL-1β expression, and differentiation of macrophages, and is required for tumor metastasis (Figure 4G).

We subsequently queried mutations, copy number alterations, and fusions of *KRAS* and *VCAN* in the cancer genome atlas (TCGA) pan-cancer dataset (*n* = 10189 patients; raw data provided in Supplemental Table 9). *VCAN* alterations (mostly missense mutations) occur in 5% of all cancer patients and significantly coincide with *KRAS* mutations (Supplemental Figure 8,A-B). *VCAN* changes show no mutational hotspot and occur in highly genomically altered and aggressive tumors (Supplemental Figure 8,C-E). In line with our findings, *VCAN* and/or *KRAS* altered cancers are the ones that commonly cause MPE (Supplemental Figure 8F). Since VCAN is a known TLR2 ligand (20), the proinflammatory loop proposed here was targeted with the TLR1/2 inhibitor Cu-CPT22 (31). The drug effectively inhibited VCAN-induced NF-κB activation in macrophages and tumor growth *in vitro* and *in vivo* at clinically relevant concentrations (Supplemental Figure 8,G-I). Hence VCAN is a clinically relevant and druggable target against human cancers that can cause pleural metastasis and effusions.

Collectively, our results show that tumor-secreted versican causes IKKβ activation in macrophages to foster metastasis in an IL-1β-dependent fashion (Figure 4G). The data from murine and human cancers indicate that *KRAS*-mutant tumors are especially prone to initiating this proinflammatory circuitry with myeloid cells, notwithstanding cancers with other mutations that are difficult to model in the mouse. Versican was previously found to activate macrophages through TLR2/6 to stimulate *Tnf* and *Il6* secretion and promote metastasis (20). We identified versican using a different approach and build upon the aforementioned study in several ways. We used bioluminescent reporter mice and macrophages in combination with an array of tumor cell lines to monitor tumor-associated NF-κB activation and inflammation and identified versican as the cardinal tumor-secreted mediator triggering NF-κB-driven transcription in host cells. We localized this response to macrophages, and employed multiple RNA- and protein-based screens to pin versican. We also identify mutant *KRAS* as an important culprit of tumor cells responsible for this inflammatory tumor-host circuit. Furthermore, we show the clinical relevance of versican and the proposed inflammatory loop in human cancers, and identify TLR1-2/IKKβ signaling in macrophages as versican’s partner in crime.

The proinflammatory interplay between versican in tumor cells and IKKβ in macrophages described here is also promising for innovations in cancer therapy and diagnosis. Multiple studies implicate IL-1β as a prime tumor promoter secreted in the cancer niche by myeloid cells (3, 7, 8, 12). The clinical importance of these findings is portrayed by the findings of the Canakinumab Antiinflammatory Thrombosis Outcome Study (CANTOS), where administration of the IL-1β-neutralizing antibody canakinumab decreased overall and lung cancer mortality by 51% and 77%, respectively (9). We show versican to be responsible for secretion of IL-1β by macrophages in the metastatic niches of *KRAS*-mutant cancers. The results position these cancers as favorite candidates for anti-IL-1β therapy, and versican as a therapeutic target in this tumor category that comprises 30% of human cancers (2, 3), alone or in combination with anti-IL-1β agents.

In addition, since early diagnosis of metastasis is key to effective cancer therapy (1, 20), versican can serve as a biomarker of metastasis. This might be achieved by monitoring local or systemic versican levels in patients at risk, or by using our NF-κB-reporter macrophages as a diagnostic platform. Indeed, our data indicate that the latter can accurately discriminate pleural metastasis from other pleural inflammatory processes, highlighting the clinical relevance of our findings.

## Methods

Methods are described in the Supplemental Methods.

### Study approval

Animal experiments were approved by the Veterinary Administration of the Prefecture of Western Greece. GLAD and MAPED clinical trials were approved by the Ludwig-Maximilians-University Munich Ethics Committee and the University of Patras Ethics Committee, respectively.

